# Canalization and competition: the cornerstone of genetic network’s dynamic stability and evolution

**DOI:** 10.1101/2024.05.13.594036

**Authors:** Yuxiang Yao, Zi-Gang Huang, Duanqing Pei

## Abstract

Grasping the fundamental dynamic property is a crucial approach for understanding living systems. Here we conduct a comprehensive study into the relationship between regulatory modes and dynamic features of gene networks. Our findings indicate that conditional constraints and competition, corresponding to canalizing and threshold regulating modes respectively, play pivotal roles in driving gene networks towards criticality. Particularly, they effectively rescue biosystems from disordered area as source of evolutionary driving force. By employing variant Kauffman models, order parameters, and stability analysis, we provide sufficient numerical evidence demonstrating the diverse and distinctive capabilities of regulatory modes in stabilizing systems. Our findings give the most systematic analysis to date on the dynamic atlas of regulatory modes, offering a framework-independent proof of genetic networks operating at the edge of chaos with evolutionary implications. Furthermore, we discus the bridge between criticality and canalizing/threshold regulating modes and propose a reasonable scheme for generating model.

## 1 Introduction

Identifying the dynamic essence of living systems constitutes a pivotal issue in the field of system biology [1]. As highly self-organizing systems, gene networks exhibit a remarkable resilience to perturbations, including signal stimulus, environmental fluctuations, and genetic mutation [2, 3]. The robustness arises from the proper orchestration of gene function modules to achieve biological adaptation [4, 5]. This relies on the joint action of components and interactions, corresponding to genes and regulatory modes, respectively [6, 7]. Numerous biological processes exhibit certain regulatory behaviors.

Conditional constraints serve as the guiding principles for determining cell fate decisions, such as development, tumorigenesis, and somatic cell reprogramming. The occurrence of specific subsequent processes relies on prerequisites. For example, properly modifying DNA methylation in zygotes ensures the normal embryonic development of mice and humans by regulating chromatin accessibility from either the paternal or maternal genome [8–10]. p53 governs cellular fate, either by rescuing from DNA damage or by activating apoptosis, acting as a gatekeeper of suppressing tumors [11]. The switch of ordered and deterministic changes in the cancer genome only occurs upon loss of p53’s normal function [12]. Reprogramming somatic cells into pluripotent cells requires the removal of numerous epigenetic barriers, such as H3K9 demethylation and opening heterochromatin [13–15].

Competition, or rather equilibrium, represents another fundamental aspect of regulatory mechanisms. A sequence of binary branches consists of mammalian embryonic developmental tree. Each point of fate divergence is precisely governed by a set of contradictory factors [16]. The predominant part at the branch initiates subsequent cascades, such as Oct4/Cdx2 during early embryonic stages leading to the formation of inner cell mass (ICM) and trophectoderm, and Nanog/GATA6 during embryo ICM stage directing the development of epiblast/primitive endoderm. The processes of cell differentiation and reprogramming also exhibit seemingly contrasting dynamics. Any bias of the relevant factors can results in differentiatial cell fates, while establishing a balance between two genetic clusters can recover the cellular pluripotency [17]. From a macroscopic perspective, it can be viewed as the confrontation between genetic forces of differentiation and pluripotency [18]. At a micro level, successful or failure recovery of pluripotency is determined by effective factor combinations [19]. Moreover, the above and other biological processes are typically facilitated by another pivotal reversible transition, epithelial-mesenchymal or mesenchymal-epithelial transition [20]. It depends on the net result of the two genetic factors, resulting in nearly binary-like opposition features [21].

The phenotypic patterns of living systems also arise from the response of regulatory signals. The autonomous formation and regulatory mechanism of zebrafish patterns are driven by a reaction-diffusion process involving short- and long-range feedback [22, 23]. The formation of body segments along with the anterior-posterior axis is facilitated by concentration gradients of growth factors. The induced self-sustaining oscillations also necessitates the coordination between global signaling and local states [24–26]. The hidden mechanism, known as monotonicity, refers to the well-organized set of positive and negative consistent regulations that form sustaining or oscillating signal patterns [27]. In a broad sense, monotonicity is homologous to competition in terms of dynamic, thereby phenotypic patterns can be viewed as additional effect of competitive regulations.

The two primary regulatory modes represent the orderliness of genetic networks in contrast to random systems, generally causing a variety of particular dynamic features. Notably, genetic networks are always at the critical regime [28, 29], enabling living systems to balance robustness and evolutionary potential [30]. These features show a strong dependency on the conditional constraint modes. On the other hand, some properties of genetic networks, such as cellular phenotypic plasticity or minimal frustration of network [31, 32], emphasize the characteristic of “competition” between two sets of antagonistic factors. These features indicate that cell fates are ultimately determined along phenotypic trajectories dictated by the “winner” genetic cluster. However, aforementioned features are mainly summarized from ordered measures, while their dynamic patterns are rarely analyzed comprehensively. What’s worse, properties are derived from dissimilar dynamic frameworks. One visible question is what are the latent relations between frameworks and their corresponding derived ordered rules. Are the features universal across different frameworks? Additionally, what are the variations of the underlying dynamics that characterize gene networks? Existing studies on gene networks mainly focus on a single variant of a special regulatory mode [33, 34]. It needs to explore how to understand the effect of regulatory modes on evolution. To our knowledge, the dynamic properties and behaviors of various types of regulatory modes have not been previously addressed. The analysis and comparisons of dynamic patterns and biosystem features under different dynamic frameworks are limited in existing studies.

To answer above questions, we employ Boolean dynamic to abstract genetic networks and regulatory modes as Boolean networks and special Boolean functions respectively. Firstly, we identify a variety of special regulatory modes included in gene networks. The results present their significant abundance compared with random configurations. Next, diverse dynamic frameworks are employed to calculate the dynamic features of biosystems, such as criticality. Not all so-called universal laws hold true across different modeling contexts. Only “criticality” emerges as a consistent attribute irrespective of the dynamic framework employed. Then *in silico* evolution simulations are used to numerically demonstrate that one of primary driving forces in evolution is establishing functional and purposive interacting relations. Further analysis of system attractor reveals the diversified dynamic discrepancy within gene networks are “at the edge of chaos”. Finally, through means of systematic perturbation analysis and percolation experiments, we point out the distinct capabilities and characteristic dynamic patterns realized by these special regulatory modes. Overall, our work provides a comprehensive understanding of genetic networks from dynamic perspective, numerically revealing the origin of criticality and a possible fundamental component of evolutionary forces. The atlas of regulatory patterns also gives guidance for constructing artificial functional genetic networks and presents potential solutions of expanded approach to establishing abstract network models.

## 2 Results

### 2.1 Overview of Boolean network and special regulatory modes

Kauffman proposed a Boolean-network-based research paradigm that captures abstract relations of genetic networks [35]. This binary representation of dynamic systems provides a an alternative modeling strategy through an abstract way. In this study, the Boolean dynamic system is selected as our primary analysis approach. Here we briefly review the Kauffman model and introduce relative concepts. Readers familiar with these backgrounds can skip this part.

The model treats gene states as binary variables, where 1 and 0 symbolize gene expression or silence respectively. Various genetic regulatory conditions for a target gene are condensed as Boolean functions (BFs), *f*. The role of a BF is to match all possible input vectors with preassigned results. Thus the state of a gene only depends on its input and configured BF as,

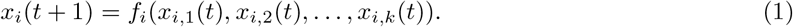

Where means that the state of gene *i* at next time is determined by the current states of its *k* input genes and *f*. If considering all genes together, one can obtain a directed network, nodes and edges of which represent genes and abstracted interactions respectively. In Kauffman’s original settings, all genes have *k* stochastic and non-duplicate inputs that are chosen from all *N* genes in the system. Therefore, the in-degree and out-degree distribution of genes are fixed value *k* and Poisson distribution with ⟨*k*⟩, respectively. Another parameter is the bias “*p*”, which represents the expected probability of “1” returned by all *f*. Under the synchronous rule, all genes simultaneously update their states. Thus one canonical Kauffman model, 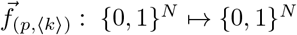, is completely deterministic.

The orderliness of classical Kauffman models is commonly measured by 2*p*(1−*p*) ⟨*k*⟩, where the value smaller/larger than 1 indicates a system is ordered/disordered [36, 37]. While this statistical measure is merely applicable to random systems and mean field conditions. These assumptions have a deviation when considering functional networks such as gene regulatory networks. One important reason is that various special regulatory modes introduce local heterogeneity and break above assumptions. Notably, these characteristics exactly arise from numerous genetic regulating modes including conditional constraints and competition. Correspondingly, some special Boolean functions can appropriately represent such modes, and we briefly introduce them as follows.

Canalizing functions (CFs) are ubiquitous components in genetic networks, usually rescuing systems from chaotic behavior [2]. These CFs exhibit dynamic patterns characterized by conditional constraints, resembling an “if-then-else” style of computer programming known as “canalization” (Figure 1a). Its formulaic presentation is,

**Figure 1:**
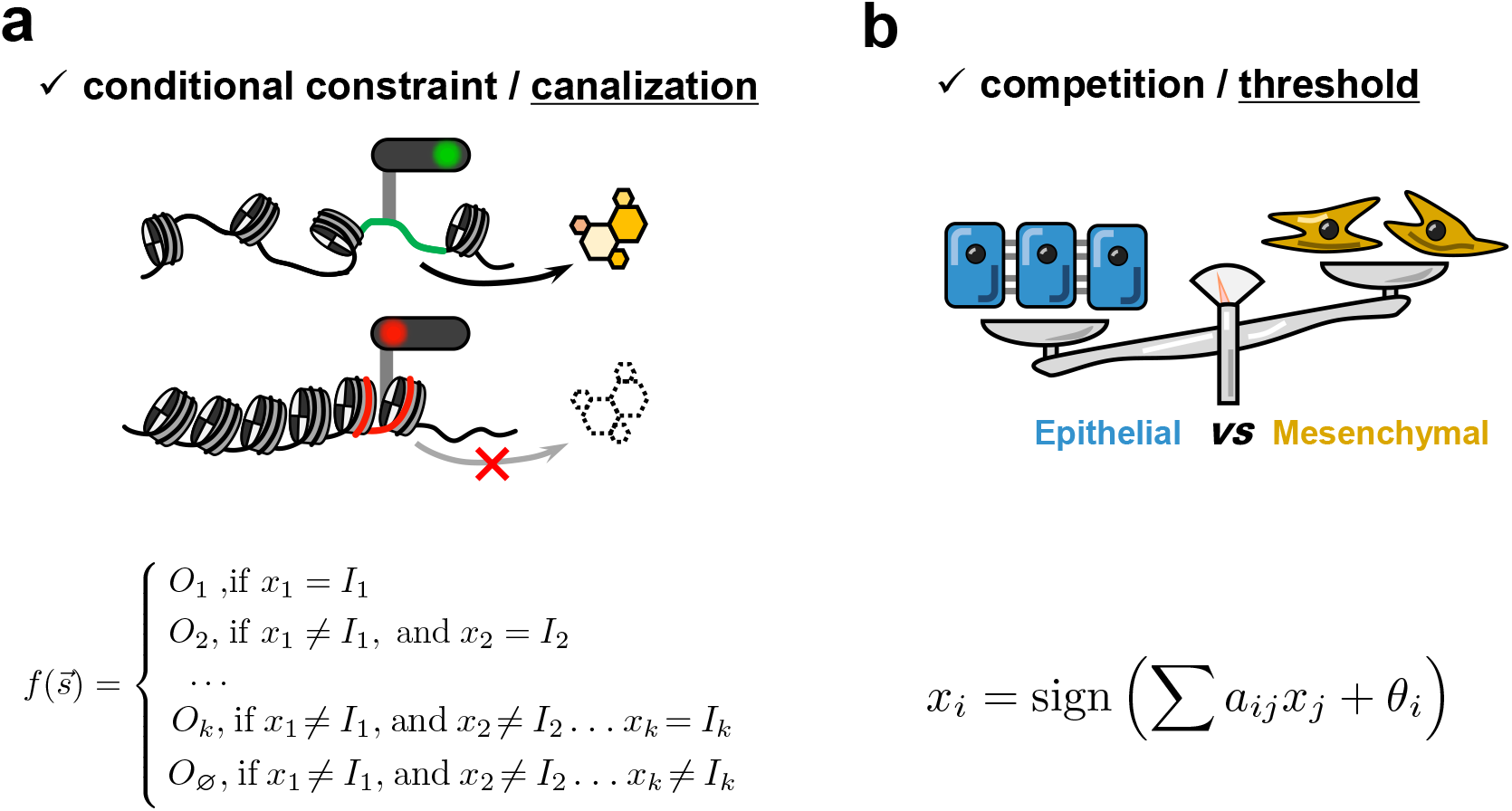
Schematic representation of special regulatory modes in biosystems. (a) Conditional constraint. During the cell fate decision process, the expression of specific target genes is contingent upon meeting corresponding epigenetic conditions, such as chromatin accessibility [13]. (b) Epithelial-mesenchymal transition and mesenchymal-epithelial transition play crucial roles in various physiological processes. The net effect of two cluster of genetic factors associated with epithelial or mesenchymal states determines the outcome[21]. These two special regulatory modes can be abstracted as canalization and thresholding.

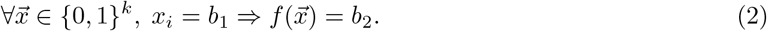

Where *x*_*i*_ means the *i*-th variable of input, and *b*_1_, *b*_2_ ∈ {0, 1}. BFs with multiple canalizing variables (*x*_*i*_) hierarchically returning series canalized values (*x*_*j*_) are called as *nested* CFs.

Threshold functions (TFs) are another representative of genetic network dynamic [38]. Each component of input contributes to the target state through positive or negative regulating effects. The final output of the targeted gene depends on their net values as (Figure 1b),

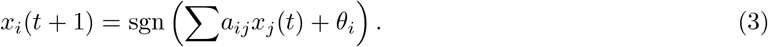

Where sgn(), *a*_*ij*_ and *θ*_*i*_ are the sign function, interaction weights, and response threshold respectively. The weights quantify the impact of source genes on the target. Thresholds represent the level of difficulty in remaining basic expression of targeted genes, for instance, *x*_*i*_(*t* + 1) = sgn (θ_*i*_) when there is no input.

Monotonic functions (MFs) are initially proposed to analyze fundamental properties of Boolean system [39]. MFs stress the consistent relation of binary vector “predecessor-successor” pairs. For example, (010), (100) are predecessors of (110), and the latter is the successor of the formers. Their mapping results should meet a sequential pattern as,

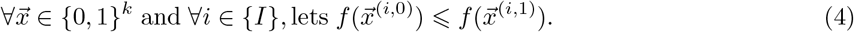

Where {*I*} is {1, 2, …, *k*}, and 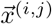 means *x*_*i*_ = *j* in the vector 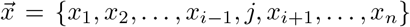. A BF that satisfies above constraint condition is called as an increasing MF. Inversely, a decreasing MF requires the “⩾” relation in Eq. (4). In biological applications, one gene can be controlled by both activatory and inhibitory regulations simultaneously. A variant of MFs with mixed constraints allows the combination of increasing and decreasing patterns [40].

For comparison, we also introduce several other special modes to explore their applications in biological contexts and dynamic properties. Effective functions (EFs) indicate that all variables of a BF are valid for regulating the output at least one case [41]. Namely, all regulations of a genetic network should be functional and effective. Formally, a *k*-input BF belonging to EF class should satisfy that,

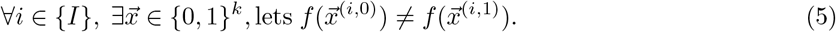

{I} and 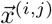 are same as the previous definitions.

Post class functions (PFs) can stabilize Boolean systems with better closure properties and consist of a larger set than CFs [42, 43]. Compared with CFs, PFs seem to keep complementary constraints. Typically, a subset of Post class *A*^*µ*^ is the functions that meet,

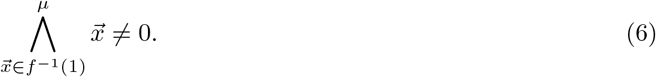

Where Λ means bit-wise AND operation, and “*µ*” indicates *any µ* 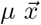 from the set 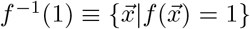 should meet the above requirements. *a*^µ^ represents the opposite case,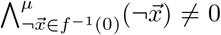. PFs indicate that all conditions mapping to a specific values should possess common components.

Dominant functions (DFs) reflect an information game mode that always returns the dominant value of the input,

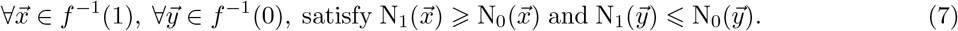

Where *f*^−1^() has the same meaning as PF’s. Symbols 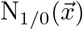 return the number of 1/0 in 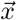. The most significant feature of DFs is recognizing and transferring the dominant value. Now we introduce all special BF (SBF) classes and represent their BF sets by using initials. The complete set can be written as 𝕊 _+_≡ 𝕊 ∪ 𝔼, where 𝕊 ≡ ℂ ∪ 𝕋 ∪ 𝕄 ∪ ℙ ∪ 𝔻. 𝔼 is excluded due to its trivial dynamic properties. Detail settings and configurations can be found in the Methods section.

### 2.2 SBFs in GeNets are abundant across dynamic frameworks

To understand the significance of SBFs involved in gene networks (GeNets), we examine their abundance contained in real GeNets. We identify the logical-based GeNets from cell collective website and categorize them accordingly. Figure 2a gives the abundance of each type of SBF. Obviously, a substantial proportion of biological BFs are “canalizing” and “effective”, which is consistent with the previous findings [2, 41, 44]. Results seem to suggest that canalization plays the role of GeNets regulatory modes. Furthermore, the fractions of BFs belonging to 𝕄 _*e*_, ℙ, 𝕋 _*e*_ are also high, with almost all exceeding 50% at different *ks*. The symbol 𝕄 _*e*_ and 𝕋 _*e*_ indicate non-strictly constraint forms of 𝕄 and 𝕋, which are suitable for more general situations (See Methods section). Inversely, BFs from 𝔻, 𝕄, and 𝕋 are relatively rare, only observed at small *k* cases. It suggests that these SBFs may not constitute the component of GeNets.

**Figure 2:**
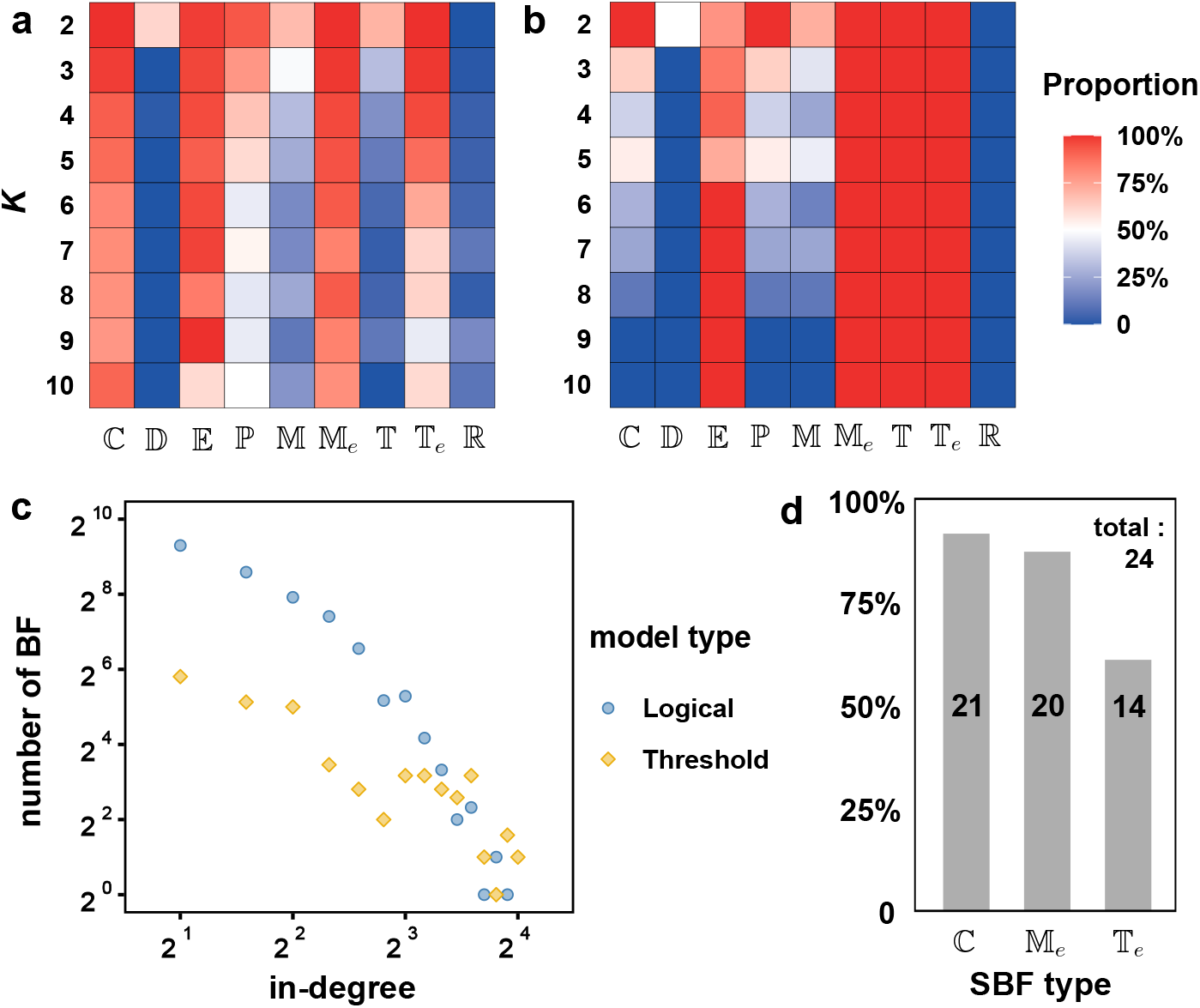
Abundance of SBFs in biological networks. Each BF of networks are categorized based on their truth tables under varying ks. Horizontal and vertical axes denote categories and k, and the colorbar represents corresponding proportions of each configuration. Logical models (a) and threshold based models (b) are analyzed respectively. (c) The in-degree distribution of vertices of all models are presented on logarithmic axes. (d) Genes with large in-degree are mostly canalized, monotonic, and thresholding. Numbers represent cases of BFs belonging to each type. Statistics of (d) only contains logical models.

To ensure the consistency of “canalization” across dynamic framework, we use another seven threshold-based GeNets and reanalyze their BF categories. According to regulatory relations, it is convenient to switch BFs from threshold-based to logical-based ones. Here we only present the results derived from standard Boolean threshold rules (variables from {0, 1}), while so-called “spin-like” rules (variables from {−1, +1}) yield identical results (SI, figure S1). All BFs certainly belong to 𝕋 regardless of their binary or spin-like forms (Figure 2b). BFs are also naturally effective due to each variable involved in inputting summation. More importantly, because of inherent features of MFs, their positive (negative) regulations definitively result in increasing (decreasing) truth table patterns, remaining consistent with unweighted TFs. 𝕋 and 𝕄 represent the distinctive feature of threshold models. However, the canalizing pattern no longer occupies a central position within the dynamic system, and only small proportions exist in cases where *k* ⩽ 5.

Even in discrete systems, the scaling law can still be regraded as another essential feature (Figure 2c). Inputs of BFs with fewer variables usually satisfy various identification criteria of SBF [45]. So we check the SBFs exhibiting high in-degree (> 9) within logical-based models. The majority of them show the features of canalization, along with monotonic and threshold modes (Figure 2d), in sharp contrast to random BF at the same *k*. Combining these findings, we conclude that the two regulatory modes ℂ and 𝕋 (𝕄 homologous to 𝕋) widely participate in establishing GeNets. Additionally, the abundance of the three?SBFs shows a strong dynamic framework-dependent effect. This highlights the importance of carefully?handling and drawing conclusions about GeNets using distinct frameworks.

### 2.3 Canalization and thresholding establish GeNet’s criticality and drive their evolution

To validate the “criticality” features of GeNets, we utilize the system expected sensitivity ⟨*s*⟩ as the metric. Sensitivity quantifies how significantly one BF is affected by variations in inputs (Definition see Methods section). So ⟨*s*⟩ represents the average sensitivity of all BFs within the system. This value exceeding one or not suggests whether the system can expand minute perturbation throughout whole system. Thus ⟨*s*⟩ = 1 reflects states that are situated “at the edge of chaos” states, where systems exist in a delicate balance of perturbation amplifying or vanishing. ⟨*s*⟩ of all GeNets fluctuate around the critical value *s*_*c*_ = 1 regardless of dynamic frameworks (Figure 3a). Logical models perform greater robustness compared with threshold-based ones due to a deviating model with s = 1.74. In fact, this model is still more critical than the random configurations (2 ⟨*p*⟩ (1 − ⟨*p*⟩) ⟨*k*⟩ ≈ 5.096), which can be also regraded as being at the edge of chaos roughly. Generally, “criticality” is always accompanied with canalizing [29], the most obvious attached feature of logical models. Our results give an explicit and universal proof that gene networks are facially critical systems from a macroscopic perspective.

**Figure 3:**
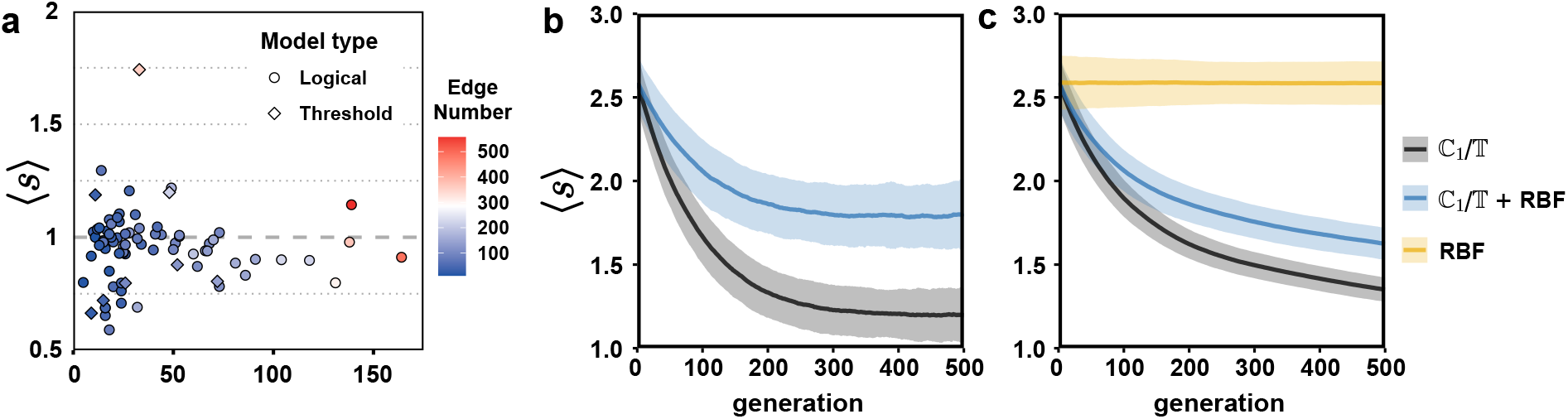
Criticality feature of dynamic framework-free GeNets and its evolutionary roles. (a) Model’s sensitivities of all models are calculated to demonstrate their criticality. Model types are presented as circles (logical) and diamonds (threshold). The colorbar indicates the number of regulatory relations contained in each bio-nets. Models are ranked based on their sizes. System sensitivity changing are simulated under different evolutionary strategies. (b) GeNets change their regulation through various evolving methods. Only difference is setting CFs+TFs at new nodes (black) or not (blue). (c) Similar evolutionary strategies are employed excluding node and edge deletions to eliminate topological influence on reducing ⟨s⟩. Blue and black curves denote whether introduce random BFs or not. Yellow curve is completely assigning nodes with random BFs as reference. Each curve is generated 10^3^ realizations for statistics. The detail changes of node, edge, and bias can be seen in SI-Fig.S2.

Despite some controversy about the criticality[30], GeNets exhibit unquestionably remarkable capabilities in realizing complex functions, adapting to diverse environments, and mitigating the effects of noise. These abilities stem from a fundamental yet apparent factor — evolution. Briefly, natural selection eliminates diverse individuals with lower fitness and adaptation, where genetic mutations play a crucial role. Here we propose mimicking the processes of GeNet evolution through various pre-designed rules. To uncover the regulatory modes caused by BFs during evolution, we observe random genetic mutation events involving gene creation/loss and changes in regulatory modes. These operations are equivalent to adding/deleting nodes or edges, swapping links between nodes, and modifying gene truth tables (See Methods).

The simulating results of random mutations indicate that natural evolution of systems leads to a decrease in complexity (Figure 3b). One contributing factor is the deletion of edges and nodes weakens coupling between genes, resulting in some isolated nodes and sparse connections. In contrast, the presence of CFs or TFs introduces a certain degree of orderliness. The difference between the two curves in figure 3b is whether newly added nodes can be only assigned with CFs or TFs. To exclude the influence of gene dysfunction resulting in node and edge deletion, we subsequently allow systems to continually expand in size and check their evolutionary tendencies. Figure 3c suggests that, beyond the topological effects, the orchestrated regulatory modes play a pivotal role in establishing order and driving the system towards a state of “criticality”. Purely random configurations maintain their complexity independent of system size and connection density (Yellow curve). The introduction of CFs or TFs through the addition of new nodes or edges, leads to a reduction in system complexity (⟨*s*⟩) compared to random configurations (Blue and black curves).

Theoretically, the structure and regulations consistently contribute to the robustness of gene networks [46]. However, robustness is not the only factor at the evolution. In certain biological cases, Park et al. suggest that stable expression of genetic interaction hubs leads to the system’s robustness. Additionally, epigenetic epistatic interactions contribute to single gene deletion phenotypes, demonstrating a canalized regulatory pattern [47]. Transposable elements (TEs) have the capacity to alter genomic architecture, induce mutations in genes, and regulate gene expression, ultimately influencing evolution [48]. Key endogenous retroelements (EREs) can exhibit conserved patterns during human genome evolution. Despite not being protein-coding genes, they significantly impact loss of function and developmental processes through interactions with species-specific proteins that recognize their sequences.

On the contrary, a morphological dynamic model demonstrates that the distributions of activator and inhibitor concentrations orchestrate various aspects of tooth development in mice and voles [49]. The differential gene activity reflects the regulatory weights and thresholds among different components, revealing competitive regulatory patterns. Analogously, experiments on yeast’s interspecies crossing demonstrate the robustness of negative-feedback regulation. Both yeasts possess evolutionarily conserved networks and only exhibit negatively regulatory effects when their promoters are swapped. These findings suggest that the strength of genetic factors, rather than condition-constrained modes, drive species differentiation.

Furthermore, *S. cerevisiae* and *S. pombe* share similar protein complexes with consistent functions. The additional components of the Set3 complex in *S. cerevisiae* highlight the redundancy of molecular regulation, as its extra orthologs, Set4. In *S. pombe*, the two orthologs of *S. cerevisiae*’s Rco1 are Cph1 and Cph2, which together form the Rpd3C complex [50]. These additional and parallel genes can be considered as temporarily invalid regulations denoted as (`new+origin`)&(…) (logical-based) or adding equally effective regulations (threshold-based). In summary, these experimental evidence illustrate canalized and threshold modes that represent conditional constraints and competitive regulatory behaviors of genes.

### 2.4 Parametric criticality masks the complexity of actual dynamic behaviors

To further explore their dynamic behaviors beyond the macroscopic effects (Figure 3), we analyze the characteristics of system’s attractors. Given that the practice in modeling biosystems [51, 52], we adopt the asynchronous rules here to assess the complexity of dynamic. Asynchronous rules cause uncertainty in the evolution paths, resulting in a state having 1 *N* possible successors. A limit cycle can manifest as either a single loop or be embedded in complex loops, unlike synchronous contexts (Figure 4a). So we identify different types of attractors and their domains as a side view of complexity. The eventual components of the system are categorized into three types: fixed points (FPs), small limit cycles (LCs), and complex state pools (CPs). A FP refers to a stable system state that remains unchanged for its any variables after a transient process. The distinction between LC and CP lies in the presence of more than 1,024 (= 2^10^) states in the complex looping pool (All states consist of a loop). These three types can represent steady states, suitable periodic behaviors, and complex functions of biosystems.

**Figure 4:**
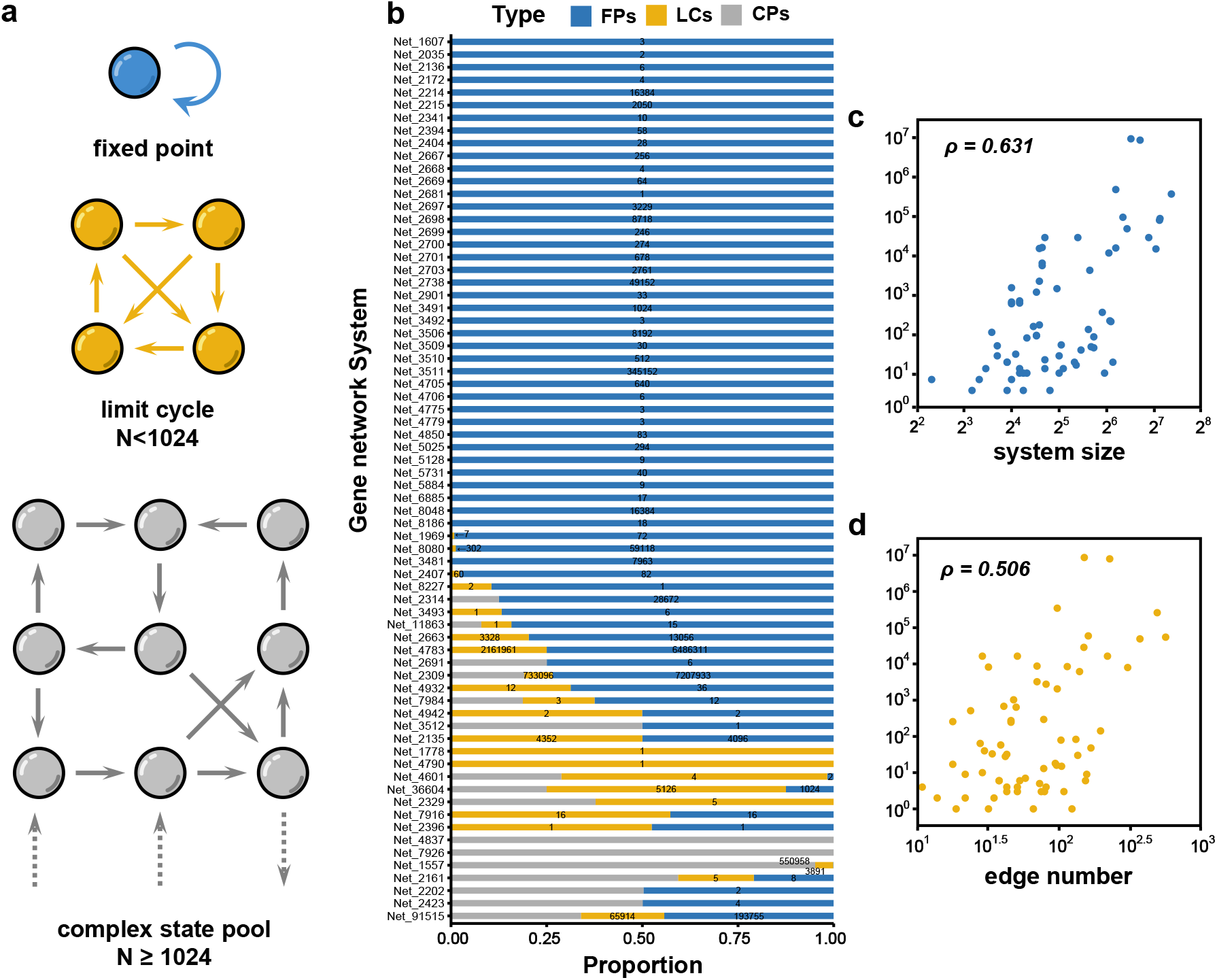
Detail distributions and numbers of attractor’s domain. (a) Schematic diagram illustrates three types of defined attractors that exists in asynchronous Boolean systems. Circles denote system states and arrows mean possible state transition directions. Note that a single state may have multiple subsequent states. (b) Real biosystems are logical-based models from Cell Collective, with model codes are labeled on the left. Horizontal axis denotes proportions of three types of attractor’s domain sizes under random sampling analyzing. Numbers above each bar (excluding CPs) represent the amounts of attractors found in simulating process. Numbers of found attractor versus system’s size (c) and connecting number/edges (d) are showed as log-log plots. *ρ* is the Pearson correlation coefficient.

We analyzed the attractor situations and corresponding domains of all logical-based GeNets. Figure 4b shows the distributions and detail numbers of the three types of attractor’s domain. Noticeably, more than half of GeNets only contain FPs, reflecting their non-periodic characteristics. Besides, their FP numbers are tremendously different, ranging from 10^0^ to 10^6^. Figure 4c, d illustrate the system size and the number of edges are partially positively correlated with the attractor number, reflecting the roles of topological links and regulatory modes. Notably, certain biosystems such as Net_8277, Net_3493, Net_7916, Net_2396 exhibit a combination of FPs and LCs, with varying proportions between them. More than half of the state space in models Net_1778, Net_4790, Net_4601, and Net_36604 finally fall in small periodic trajectories. While other FP-LC mixed biosystems like Net_1557 only show small fractions of domains. We ignore the number of CPs in the system due to the truncation of the maximum recursive threshold (See Methods section). These results suggest that critical systems can behave dramatically differently, despite their global order parameters ⟨*s*⟩ indicating a state “at the edge of chaos”. The parametric criticality dose not equate to critical dynamic.

Paradoxically, the presence of BFs belonging to ℂ and 𝕋 is common in GeNets (Figure 2). While the variety of attractors and domains seems to prompt that SBFs in restraining system consistency may not be as significant as expected. To explore their practical effects on systems, we conduct an analysis on a series of artificial GeNets generated under various evolutionary conditions. To simplify the analysis, we employ two evolving methods, changing regulations and swapping edges (See Methods section for details), which can keep the numbers of nodes and edges constant. Attractor domains belonging to the same types are entirely counted without distinguishing specific attractors. The regulatory mode (BFs) serves as the sole means to obtain orderliness. Figure 5a-c clearly shows the change in the distribution of the fraction of attractor domains for different evolution generations. As ordered dynamic regulations are gradually established, GeNets exhibit a preference for stable or small periodic attractors over complex loops. This indicates that SBFs effectively constrain regulating information as biosystems progressively obtain the “criticality”.

**Figure 5:**
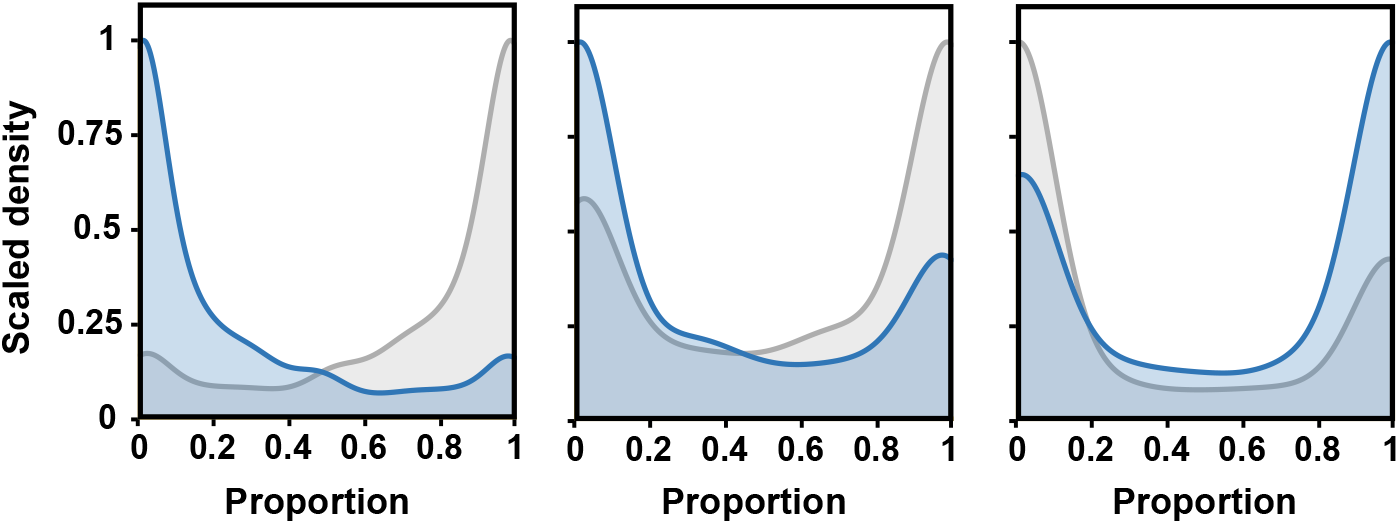
Attractors influenced by introducing CFs and TFs during the process of evolution. Distribution of two types of attractor (blue for FPs, gray for CPs) after evolving 50, 150, and 250 generations from left to right. Each evolution process is mutually independent and repeated 1,000 times. Note that vertical axices of distribution diagrams are scaled by maximum for visual consideration. LC is ignored due to its negligible proportion.

Combining that nearly all genetic BFs are canalized and thresholded (Figure 2), we conclude these two special regulatory modes play one of the central roles in establishing orderliness from chaotic-like dynamic patterns. Despite the presence of some CPs, most biosystems show much more ordered than purely random configurations (Figure 5). On another scale, one must realize that dramatic differences of system dynamic may be covered and simplified by global order parameters. A clear definition of the criticality associated with dynamic of GeNets should be proposed.

### 2.5 SBFs stabilize hybrid Kauffman models diversely

To validate SBFs in stabilizing system quantitatively and specifically, here we employ a variant form of the Kauffman model [35], hybrid Kauffman models, as the benchmark system. They consist of nodes partially assigned with random BFs and SBFs (See Methods). Previous studies indicate that 2*p*(1−*p*) ⟨*k⟨* > 1 means a slight perturbation will probably spread through whole system, called as “damage spreading” [36, 37]. From a dynamic perspective, the normal Hamming distance between one system state 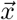 and its perturbation 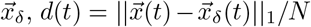, loses the stability at *d* = 0, and ultimately leads to *d*(∞)→ *d*^*^. We simulate all types of hybrid Kauffman models to determine their final Hamming distances (*d*(*t*), *t* = 1000). The black curves in figure 6 display the effectiveness of five types of SBFs in reducing system disorder. As the fraction of SBF increases, final distances decrease to lower values, approaching zero or near-zero as ℂ _2_ and ℙ display. ℂ _1_, 𝕄, and 𝕋 show less stabilizing effects than the former two. Remarkably, systems consisting of 𝔻 have aberrant intervals (Figure 6, the dotted curve), which is caused by the filtering setting of the “dominant” input. We check the distribution of final distances at the parameter proportion = 0.8 but the initial states are 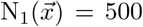 and 900 respectively (Figure 6b). The results confirm that the initial state also show a significant effect on the final distance within the dominant regulatory mode. The asymmetry of system state directly restricts the opportunity of overturning of majority and minority states. Due to the initial state’s proximity to the symmetrical points 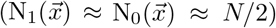, a slight perturbation can spread though the entire system, even DFs occupy a large fraction. The strong attribute of “dominant” filtering drives the perturbed state 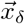 to converge towards attractors close to each other or opposite dominant states. This suggests that DFs exhibit a distinct regulatory pattern of “minority-majority” state competition, setting them apart from other logical-based BFs such as CFs. More importantly, this regulatory pattern is invalid for random BFs. To verify this hypothesis, we slightly modify the configuration of initial state 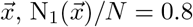, and then reanalyze the 𝔻 type hybrid model (Figure 6c). The curves of the final distance shows a sharp jump within the interval 0.6 ∼ 0.8, unlike other cases where the distance gradually varies. It implies that DFs exert a stabilizing effect on the system depending on their appropriate proportion.

**Figure 6:**
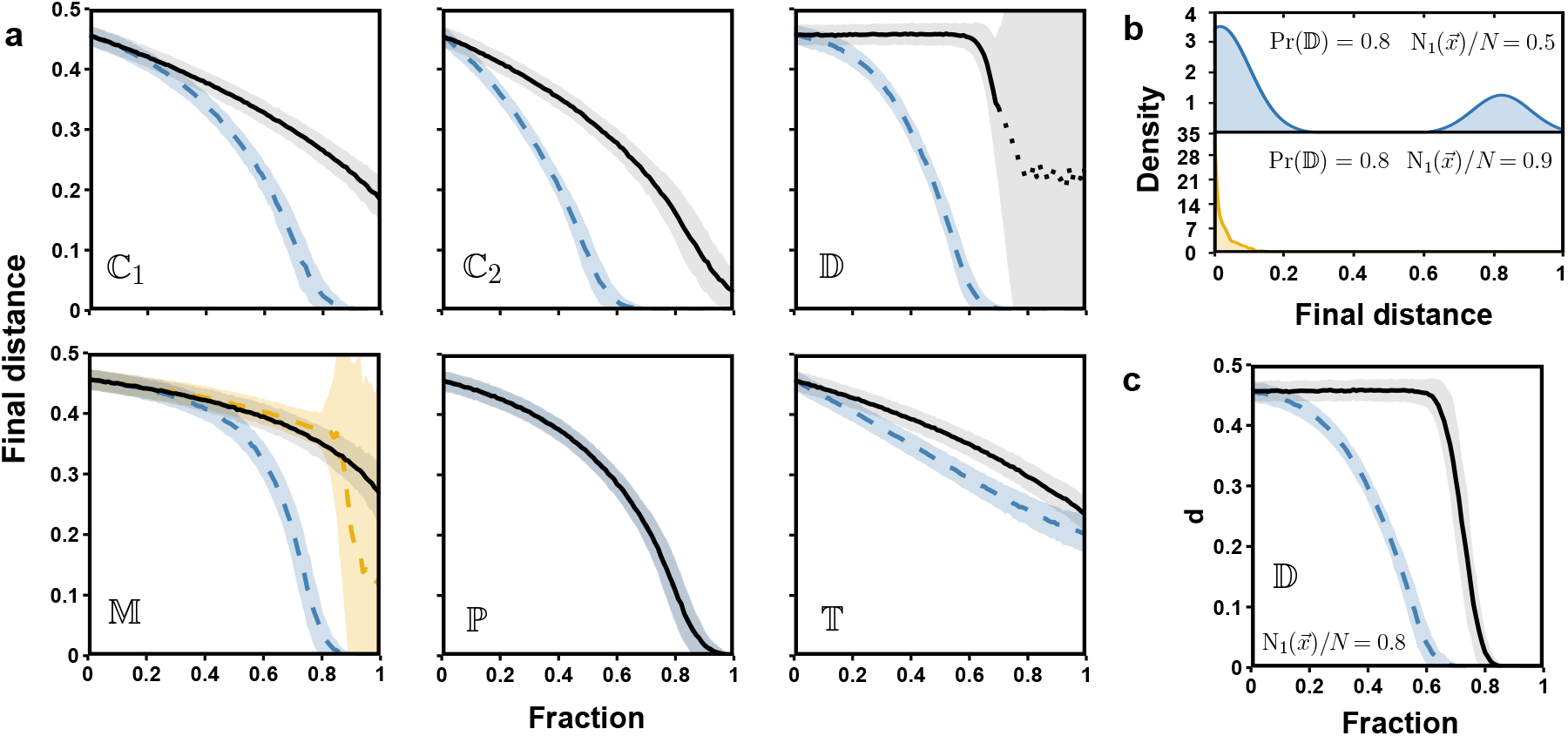
Average final Hamming distances of hybrid Kauffman model vary with the proportion of SBF mixed in system. (a) The horizontal axes and the vertical axes represent the fraction of SBFs and final distance at t = 1000 respectively. In each case, the hybrid models are generated by two methods, namely arbitrary setting and ordered constraint, depicted by black solid curves and blue dashed curves (See Methods for detail). Types are labeled in the corner, and each configuration is generated 10^3^ realizations for statistics. Dotted curve in case 𝔻 denotes the aberrant average. Blue and yellow dashed curves represent increasing / decreasing subtypes of MFs in case 𝕄. (b) Distributions of final distance of case 𝔻 under two initial configurations. (c) Power of stabilizing system of DFs can be reflected by an unbalanced initial state configuration.

To exclude the arbitrarily setting in the hybrid models, we also consider a more ordered constraint that only one subtype of specified SBFs is allowed. For instance, BFs originating from ℂ _1_ have four (2^2^) regulatory patterns, thereby canalizing and canalized values that can be freely assigned as either 0 or 1. So various subtypes of a specified type of SBFs can coexist within systems. Blue dashed curves in figure 6a,c present the corresponding final distances under ordered constraint. The ℂ, 𝔻, 𝕋 cases show that only one single subtype can effectively reduce system disorder, whereas this is not true for 𝕄 and ℙ cases. Diverse subtypes of ℙ do not introduce additional disorder. In other words, their dynamic properties of stabilizing system have no distinction among any two subtypes. However, in the case of 𝕄, monotonically increasing and decreasing MFs play inverse roles in regulating system (Blue and yellow dashed curves in figure 6a, 𝕄). One explanation is that the truth tables of these two types of SBFs have evident geometric features rather than inputting orderliness (Figure 7). The fundamental structure of a MF involves enumerating all pathways of randomly selected ancestors to “all-one” vector 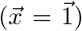. In regulatory circuits, monotonically increasing MFs typically maintain consistent patterns, serving as the role of delivering signal. Inversely, monotonically decreasing MFs tend to have a “contradiction” tendency, resulting in longer attractors in systems [53] and consequently introducing greater risks of disorder. Not strictly speaking, MFs with increasing/decreasing patterns can be analogous to multi-input buffer/inverter gates. While PFs stress common components among the selected vectors. Generating a PF is a process of choosing isolated vertices, edges, or other interconnected components (triangles, tetrahedras, …) mapping to one (*A*^2^) or zero (*a*^2^) [54]. There is no specific variable (canalizing variables in CFs) or binary value (majority or minority in DFs) that definitively modulates the flow of input-output information. We call this dynamic feature as *unbiased nature of orderliness*.

**Figure 7:**
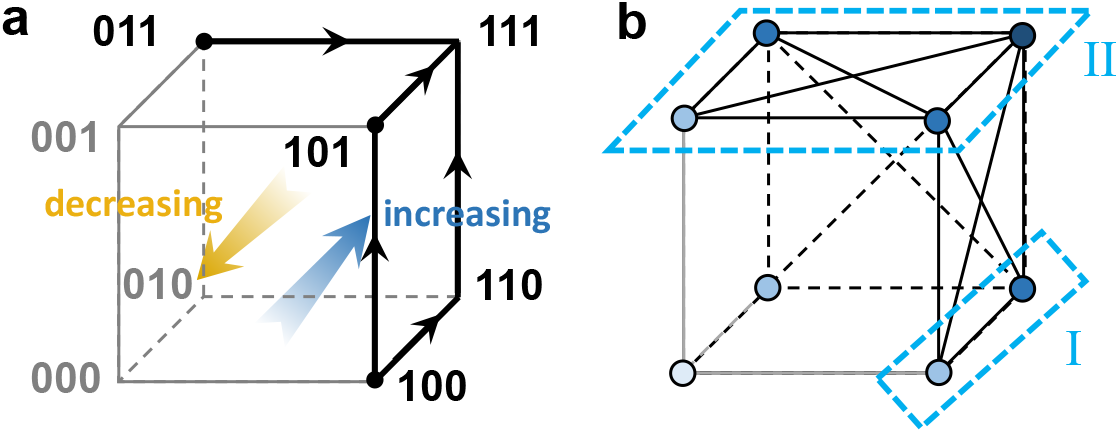
Schematic diagrams of a possible k = 3 MF (a) and PF (b). Eight binary vectors are embed in a cubical lattice, making Hamming distance of neighbor vertices is one. (a) For the MF cube, ancestors (black dots, 100, 101, and 011) and their successors are assigned to one (black vectors), while another three vertices are zero (gray vectors). Arrows indicate the pathways of increasing tendencies in the vectors. Macro-trend is vertices more likely be assigned by one from 000 to 111. (b) Vertices of PF’s cube denote same binary vectors as MF’s, and the shade of their colors represent the number of 1 included in vectors. Here we consider subclass *A*^2^, and black solid/dashed edges are candidate edges. Light blue dotted borders show two possible configurations of *k* = 3 PF, and vertices contained in the borders map to one.

### 2.6 Percolation of models reveals SBFs establish the local order

To address spatially constrained effects and static features of SBFs, we embed hybrid models within specific geometric structures such as square lattices. The spatial constraints can alter the dynamic properties of the systems. Here we observe whether the stable components of systems can form a pass-through cluster, or say a “percolation” [43]. A stable component is defined as a cluster of vertices that remain constant over a certain time after the transient process. This approach provides an alternative perspective on capturing static features of SBFs by measuring how frequently vertices change their values.

The results reveal slight differences from spatial-free hybrid models. As figure 8 shows, when subjected to arbitrary setting (black curves), ℂ and ℙ percolate earliest, followed by 𝕋. The phase transition points of ℂ and ℙ fall within the interval 0.4∼0.6, while the probability of 𝕋 percolation improves progressively. However, 𝔻 and 𝕄 appear to have limited impact on reducing the system oscillation significantly. Their phase transition points are unclear as cases of ℂ and ℙ with less frequent percolation. It indicates that the mixed cases of 𝔻 and 𝕄 are weak in regulating the local information flow. If limited to ordered constraint, all systems further effectively reduce the oscillatory behavior (Blue curves in figure 8). Cases ℂ and 𝕋 occur percolation at smaller fractions of SBFs. The curves of case ℙ once again verify the *unbiased nature of orderliness*. The curves of arbitrary setting and ordered constraint conditions almost coincide with their performance in spatial-free hybrid models. Notably, the percolation phenomena of cases 𝔻 and 𝕄 become more frequents, and even occurring in advance. This suggests that their homogeneous configurations play a more prominent role in orchestrating information flow than the cases of ℂ, ℙ, and 𝕋.

**Figure 8:**
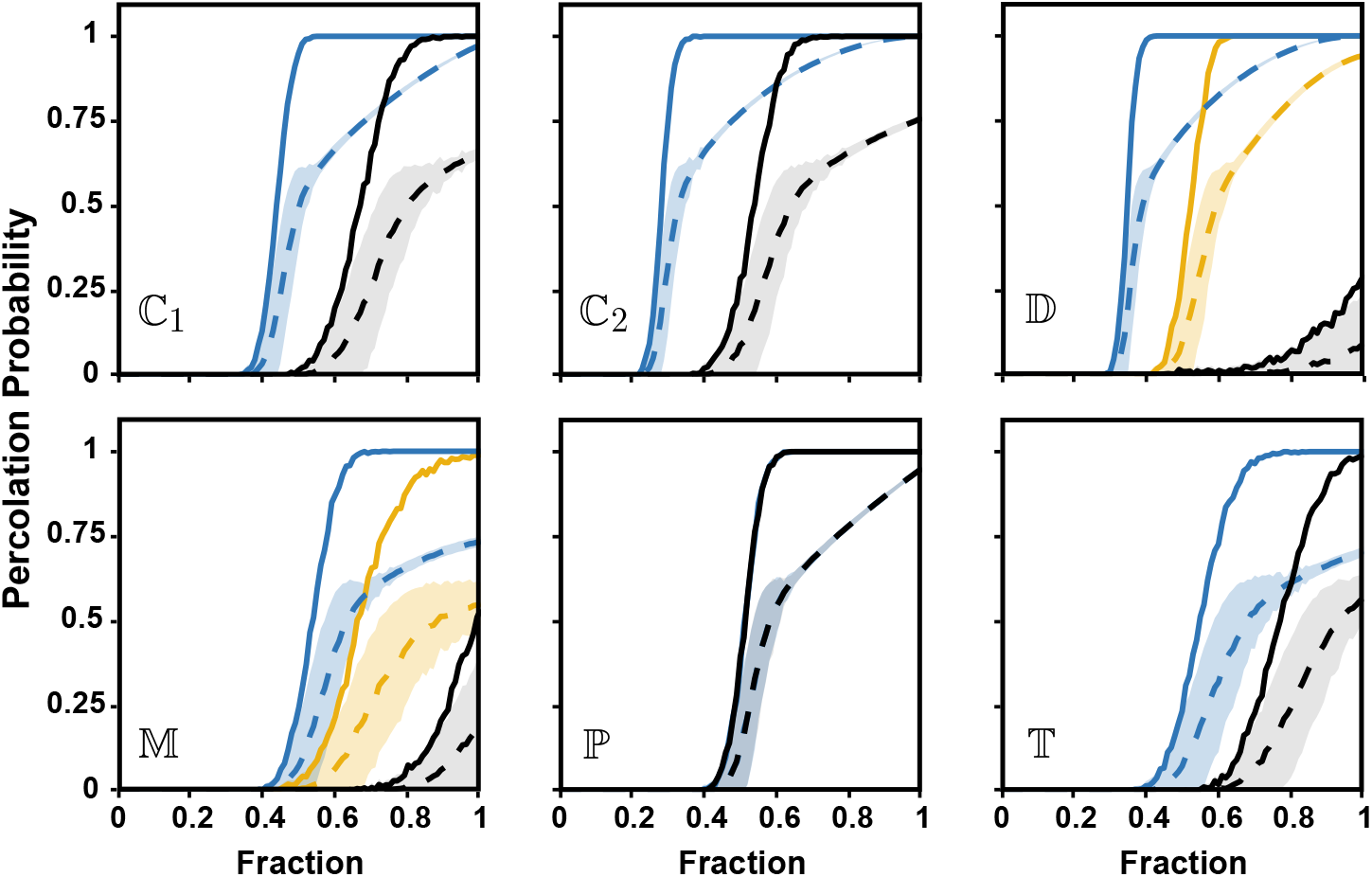
Percolation and the largest cluster of stable lattice points in embedded hybrid models. Horizontal axes represent the fraction of SBFs in the system. Vertical axes depict the proportions of percolating occurrence (solid curves) in 10^3^ realizations and the average proportion of the largest clusters of stable lattice points (dashed curves). Color blue still stands for ordered constraint compared with arbitrary setting (black). In case 𝕄, blue and yellow denote increasing/decreasing monotonicity of MFs configuration respectively. The yellow curve in case 𝔻 shows the percolation under unbalanced initial states 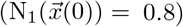, and different conditions are distinguished by color.

We also consider the special circumstances of 𝔻 and 𝕄, demonstrating the effects of unbalanced initial state configurations and types of monotonicity on stabilizing vertices. If 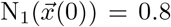, systems with 𝔻 cases successfully transform the trivial cluster into percolations at the point ∼0.5. Especially, here we do not impose any restrictions on the specific type of DFs employed, whether they are one- or zero-preferred functions. The dominant binary value effectively drives systems to achieve local self-consistency and stability. Besides, the local stabilization can also be observed in case 𝕄. As previously mentioned, MFs exhibit two primary patterns (Figure 7a) that can be roughly categorized as acting either as a buffer or an inverter gate. The increasing pattern reinforces the tendency of local information, on the contrary, the decreasing one enhances the value stability through a “double negation” way. Figure 9 gives examples of two small 50×50 square lattices to present distinct system state patterns after passing transient process. In the increasing case, vertices with same properties (values or stability) aggregate into homogeneous clusters. Conversely, stable components of the decreasing case form a checkerboard pattern, reflecting a “contradiction” frustration structure.

**Figure 9:**
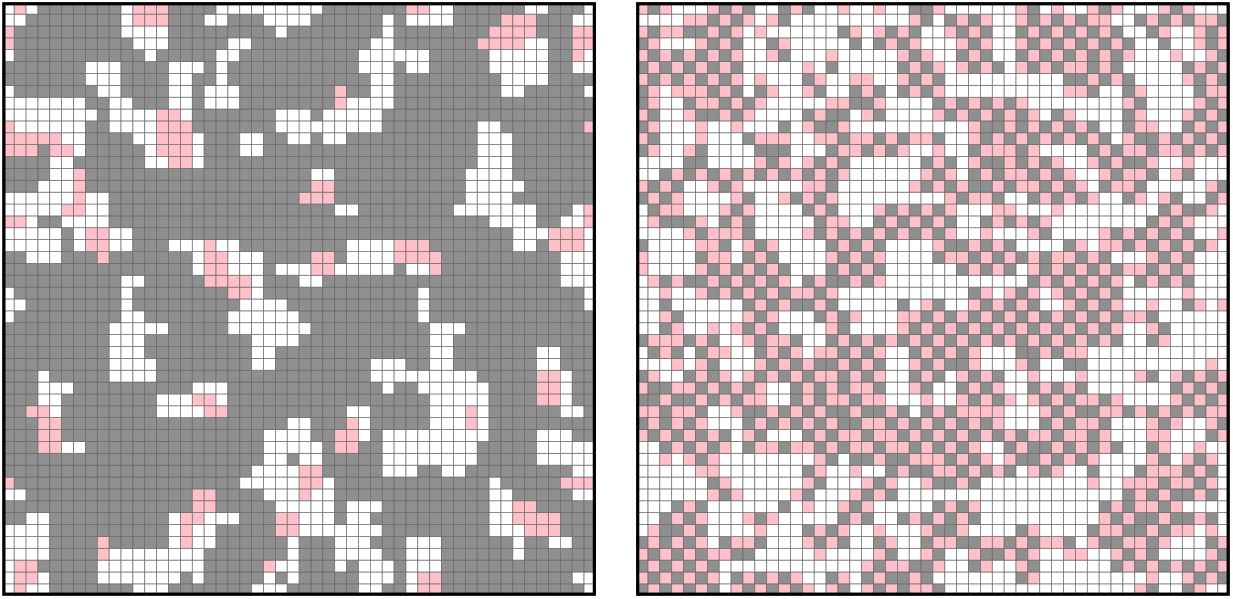
Stable state patterns of case 𝕄 with only increasing (left) and decreasing (right) MF configuration. White squares denote non-statical vertices, while stable components are colored by gray (for 1s) and pink (for 0s). Both models are exclusively consist of monotone functions and simulated from balanced random initial state, 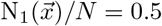.

Beyond occurrence rates of percolation, the presence of SBFs leads to varying increases in maximal clusters (Dashed curves in figure 8). The typical feature is the clusters under order constraints are larger than those set arbitrarily. Moreover, models with fractions closing to one reveal that different types of 𝕊 preferentially lead to stationary (ℂ, ℙ, 𝕋) or oscillatory (𝔻, 𝕄) attractors. These findings elucidate how?increasing numbers of CFs and TFs correspond to decreasing proportions of complex state pools, as shown?in figure 5.

### 2.7 Sensitivity and inherent order of SBFs are distinct

To further quantify the capability of SBFs in stabilizing system, we calculate the sensitivity of each subclass. This index measures how sensitive a BF mapping results are changed by its inputs,

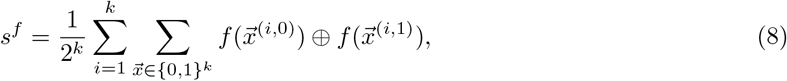

Where ⊕ is Boolean exclusive OR operation. Figure 10a shows the sensitivity expectation (⟨*s*⟩) of each subclass exhibits distinct rates of increase with respect to *k*. The indices of all subclasses, except for EFs (not presented here), have significantly smaller values compared with random configurations. This observation tentatively suggests BFs from 𝕊 can stabilize systems to varying degrees.

**Figure 10:**
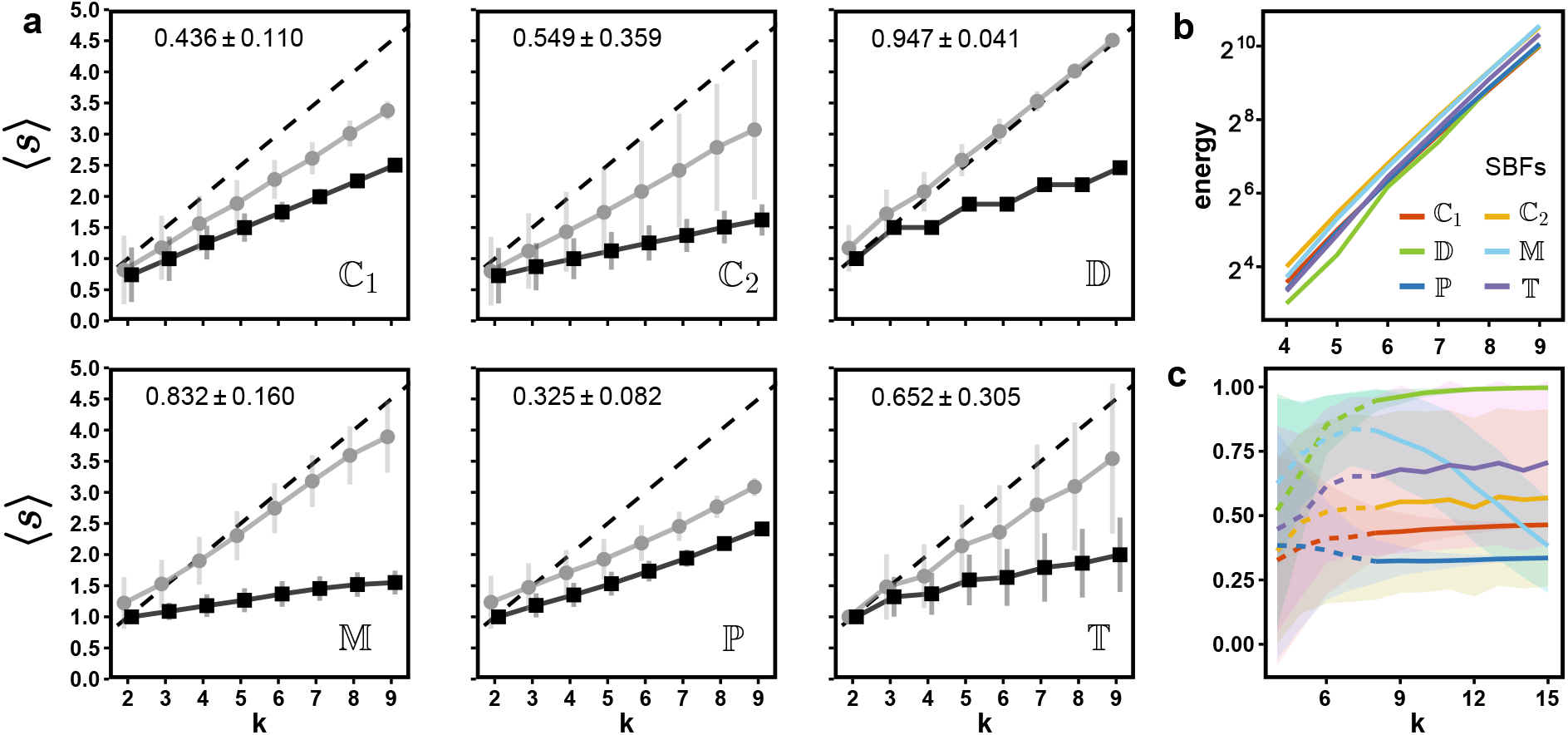
Function sensitivity and truth table pseudo energy of 𝕊. (a) ⟨*s*⟩ of SBFs (black square) and their shuffled configurations (gray circle) are plotted against *k*, generating 10^3^ realizations. Type and intrinsic bias (under the *k* = 8 case) are labeled in the corner. (b) Pseudo energy increases exponentially as *k* varies across all 𝕊. Small confidence intervals are not shown here to avoid visual interference (The separate curves in SI, FigS3). (c) Solid curves depict the relative energy of 𝕊 versus *k*, while dotted curves imply the uncertainty due to small *k* values. Colors are same as (b).

The sensitivity of each subclass has a fluctuating range with respect to the given parameters. In the cases of 𝕊 \ 𝕋, ⟨*s*⟨ tends to converge as *k* increases. The primary reason is that a larger value of *k* leads to the truth tables closer to the defined intrinsic features, thereby reducing randomness of the configuration. It also means the dynamic behavior of 𝕊 \ 𝕋 is well predictable. From a biological perspective, CFs introduce greater certainty in special input variables and fixed results (canalizing and canalized values) than TFs. Case 𝕋 shows opposite results due to the arbitrary configuration of positive and negative inputs. A larger *K* value facilitates a shift in the weighted summation away from the threshold, resulting in more ineffective inputs and less sensitivity. The delicate stability exhibited by TFs configured near the threshold, highlights their heightened responsiveness to inputs. As another side of the coin, the wide range of sensitivity also provides TFs more potential dynamic redundancy, such as evolutionary capabilities.

The numbers in figure 10a indicate corresponding *intrinsic bias*, which refers to the proportion of the majority state in truth tables regardless of specific binary value. These show the significant state biases of SBFs in their truth tables. The gray curves in figure 10a show corresponding ⟨*s*⟩ after shuffling truth tables. The shuffled cases indicate *intrinsic bias* also plays a partial role in reducing sensitivity. Actually, any slight deviation from *p* = 1/2 can effectively reduce BF disorder. To evaluate this reduction, we propose a truth table pseudo energy 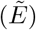 that measures the order of SBFs. 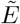 has the analogy with magnetic systems, but only focuses on local heterogeneity,

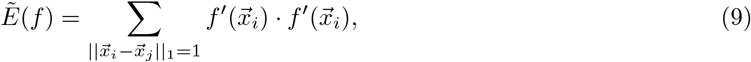

where 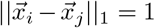 means the enumeration of each pair of input vectors with Hamming distance of 1. *f*^′^ is the spin-like substitute of the original *f*, thereby its transformation can be written as 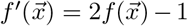. The energy describes “spin” frustration in the graph of binary hypercube lattice (SI, Figure S3). For random BFs (*f*_*r*_) with identical *k*, larger |p 1/2| corresponds to higher 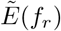. It leads to 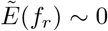 due to homogeneous distribution of “spin”, especially 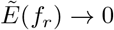 when *k* ≈ 1. We treat this case 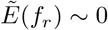 as the “ground state” of BFs, which acts as the reference of fundamental disorder. 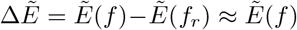 stands for the demand energy of establishing order of a BF. Part of 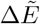 is resulted from *intrinsic bias*, denoted as 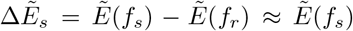, where s refers to shuffling truth table of *f*. The ratio of 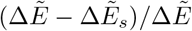, relative energy (RE), represents the *intrinsic dynamic modulating* capability of SBFs.

Figure 10b shows 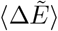 of all types of SBFs. The cost of establishing all SBFs increases exponentially with k due to length extension of truth table. Inversely, SBFs possess a diverse range of capabilities of *intrinsic dynamic modulating*. Figure 10c shows that as *k* increases, the ratios of 𝕊 approach respective constants. It is evident that 𝔻 and 𝕄 include a relatively obvious intrinsic modulation followed by 𝕋, while the *intrinsic bias* of ℂ _1_, ℂ _2_, and ℙ plays dominant roles in 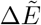. What’s more, some intervals of ⟨*RE*⟩ of SBFs are quite wide (Figure S4), suggesting that those SBFs have potential plasticity of *intrinsic dynamic modulating*.

## 3 Discussion

In this study, we analyzed special regulatory modes in biological systems, confirmed the pseudo-criticality with distinct dynamic behaviors beyond it, and comprehensively explored the foundational origin from a dynamic perspective. Primarily, the concept of canalization is proposed to describe genetic network and evolution [55], which is subsequently applied to genetic systems through Boolean networks [35]. This strong signal-constraint effect resembles an “if-else” cascade regulatory modes via making non-canalizing variables invalid when canalizing input exists. Henceforth, canalized functions, notably reducing system disorder, have long been regarded as the cornerstone of gene network stability or criticality origination, and are extensively explored in relevant studies [2, 34, 44, 56]. However, “canalization” are insufficient to encompass all biological processes. Inspired by neural networks, threshold-based models are proposed, and the have been shown to share common features with classical Boolean model [57]. It has found extensive application in the modeling of genetic networks, such as denoting cell cycle of yeast [58, 59], studying genetic network evolution [60], and elucidating cell fate decision [7, 31]. One characteristic hallmark of genetic networks is the antagonistic interplay between two clusters with opposite properties, such as epithelial-mesenchymal transition [20, 31], cellular pluripotency and differentiation states [17, 29]. There exists numerous redundancies in regulatory relationships, and their net effect drives cell towards a determined fate. This obviously contradicts the “if-else” pattern.

The monotonic modulation exhibits two prominent characteristics: tautology and contradiction, which correspond to regulations of increase and decrease respectively (Figure 7a). These two central roles both transfer and shape the signals of living systems. Figure 9 shows a tautological pattern that transmits the state flow, following a global tendency. Contradictory patterns, on the other hand, usually prefer to form local signal-state antagonisms and exhibit regular patterns. This mechanism assumes a significant role in biological processes. For example, during human segmentation development, antagonistic gene pairs associated with WNT, FGF, and Notch pathways collectively contribute to the formation of paraxial mesoderm segmentation and the establishment of segmental clock [25, 26]. Signals within this pathway are crucial for maintaining an overall regulatory trend, which holds significant biological implications in terms of monotonic modulation. Here we speculate the emergence of checkerboard patterns at critical intervals can be attributed to the suitable monotonic configurations [61], after all thresholding and monotony share same dynamic rules as discussed above.

Genetic mutation plays a pivotal role in driving the diversification of biosystems, contributing to fitness and adaptation that is the measure of species evolution. In this study, we focus on regulatory modes, special Boolean functions, while ignore the topological effect on the system orderliness. By simple verification *in silico* (Figure 2-5), we reveal that conditional and competition dynamic behaviors are common in genetic networks. In particular, the numeric evidence obtained from evolution simulations shows CFs and TFs can rescue biosystems from falling into chaos. To be sure, our mutating strategies neglect many details, such as mutation rate, gene interaction differentiation, which need to be integrated with more evolutionary biology knowledge. Apart from that, an implicit assumption we used here is that FPs or LCs can play more biologically meaningful roles than CPs. A biological process should not pass through too long or too complex cellular states; however, the presence of complexity in dynamics also preserves the potential capability of adapting to dramatically changing environments. This equilibrium may offer an alternative interpretation of the so-called “criticality”. Although certain limitations in the analysis, the various distributions of attractor domains reveal neglected attractor preferences behind the nearly perfect criticality index. Nevertheless, this distinction in domains also implies a potential approach to roughly evaluate the degree of evolution. For a common subsystem across species, checking their attractor types and domains can give a tentative measure of the species position on the evolutionary tree or indicating key genetic events, such as metabolic preference and DNA transposon accumulation [62].

For dynamic behaviors, when mixed with arbitrary fraction and type of SBF, systems undoubtedly reduce their disorder more or less, thereby approaching a state closer to “criticality”. It should be noted that 𝔼 is a result-oriented feature without any priori order constraints. Strictly speaking, it cannot be regarded as dynamic special modes, one can ignore this type BF in modeling genetic network. BFs from 𝔻 are employed to illustrate whether the majority/minority competition is appropriate for genetic networks, as they are used in social modeling [63]. The answer is negative. However, DFs are a subset of TFs that meet *A*_*ij*_ = 1 and *θ* = *k/*2, which results in small *intrinsic bias* but the worst capability of stabilizing system (Figure 6, 7, 10a). Monotonicity is one of crucial concepts of BFs [39]. In a dynamic context, it shows the tendency of regulating patterns from upstream to downstream. As shown in figure 2b, monotonicity is actually associated with the threshold-based Boolean functions, whose dynamic basis are consisted of weights and thresholds. The positive/negative weights contributing to the summation remain consistent, inevitably leading to the *generalized* monotonicity. In a broad sense, 𝕄 is homologous to 𝕋. It must be noted that our monotonicity refers to the pattern of variables (𝕄_e_), not the global tendency (𝕄, see Methods). One special type is the Post class, whose subclasses show the homogeneity of dynamic patterns (Figure 6, 7). The reason is that the two subclasses (*A*^2^, *A*^2^) exhibit symmetry through 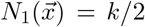 hyperplanes, resulting in identical truth table structures and dynamic effects. Despite their outstanding ability to stabilize systems, PFs are relatively uncommon in real biological regulatory systems (Figure 2). In this way, the typical features of biological systems are hidden in ℂ and 𝕋.

By using the sensitivity of Boolean functions, we quantified their susceptibility of changing variables. Another order measure is calculated from the pseudo-energy index. The “energy” depicts the cost of establishing the order of truth tables. Intuitively, the order of SBFs arises from intrinsic bias and intrinsic dynamic modulation, both of which constrain the SBF outputs and lead to ordered dynamic patterns. From a system perspective, we evaluated the damage spread of small perturbation across various hybrid Kauffman models. Our results show BFs from ℂ and ℙ can effectively reduce the system disorder. MFs exhibit distinct performance in terms of monotonicity (increase/decrease), TFs offer a wide adjustable range, while DFs strongly relying on the initial state distribution, reflecting SBF’s diverse signal modulation behaviors. The results of stable component percolation also confirm above conclusions. The percolation of hybrid models with ℂ and ℙ occurs earlier than other types, followed by cases 𝕄 and 𝕋. The homogeneous type of a specific SBFs can significantly enhance system stability compared with general cases (Figure 6 and 7, blue curves).

In a word, ℂ and 𝕋 both exhibit the ability to mitigate system disorder and possess distinct biological significance. Therefore, beyond ingenious topological structures and appropriate function assignments [6], the two types of SBFs occupy the central status of modulating information patterns and further shaping system behaviors. Nevertheless, we must recognize that behind the conventional criticality phenomena lies a system preference of diverse functions (Figure 2, 3), where logical models highlight the canalization but threshold ones underline monotonic mechanics. Considering that strong influence of canalizing function on systems, several classical GeNet features, such as minimum complexity [45], minimum-sensitivity [64], or redundancy [56], are primarily related to canalizing functions rather than the general properties of GeNets. Modification based on advances in biological findings, taking into account actual cellular systems instead of descriptive inference, can avoid our falling into the worship of canalization and the pitfall of model bias.

More generally, we conclude that a single class of SBFs is insu?icient to realize complex capabilities of genetic networks. The emergence of a highly adaptable living system results from the integration of diverse functions with appropriate topological connections. In light of this, we propose three prospects for future study. Firstly, from the evolutionary angle, it is crucial to explore the events give rise to various types of regulatory modes in biological systems and understand how these special regulations enhance the adaptability and fitness of genetic systems [30]. Next, with advances in biotechnology, we have gained a deeper understanding of biochemical components and genetic networks. It is time to reassess the canonical Boolean framework and consider the combination of canalization and threshold selection or introduce more discrete values [65]. For instance, the balance or competition of pivotal transcription factors (threshold) and the requisite epigenetic precondition (logical) are common features in cell fate control and decision-making processes [13–15]. In the end, the simplicity of Boolean networks still renders them highly vitalized, yet their integration with state-of-the-art bioinformatics algorithms and omic data is imperative to capture the essence of genetic networks.

## 4 Methods

### Special function settings

Configuration of *f ∈* 𝕊 for nodes is randomly assigned, unbiased, and independent of node degree. It means the specificity is independent of in/out-degree or other topological features.

Canalizing functions are categorized into two types: single canalized and *nested* canalized. In the case of single canalization, BF has only one canalizing (input), corresponding to one canalized (output) value as Eq. (2) shows. While the *nested* one stresses hierarchical regulating patterns,

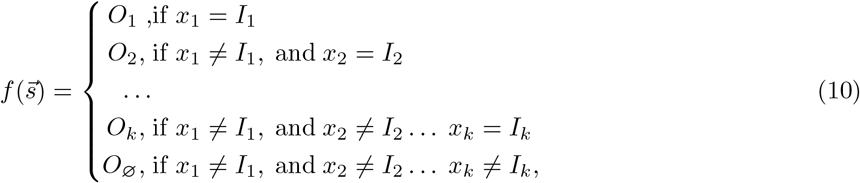

where *I* ∼, *O*∼∈ {0, 1}. Note that {*x*_1_, *x*_2_, …, *x*_*k*_}can be reshuffled or only part of them act as canalizing variables. In our analysis, we take the two-level cases (only *O*_1_ and *O*_2_) as representative of nested CFs, which implies two variables in BFs can canalize function results. The value of *I*∼, *O*∼have no limit under arbitrary settings, while the ordered constraint requires keeping a constant value.

Dominant BFs are conveniently configured to generate the truth tables by comparing 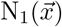 and 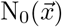 of input vector 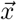,

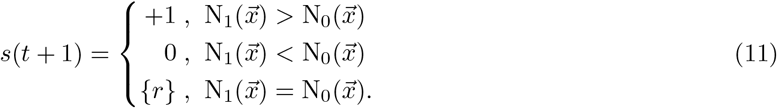

Where {*r*} means a random assignment to 1 or 0 in one function generation. The difference between ordered constraint and arbitrary setting is whether fix the “random” or not.

Effective functions constitute a large proportion of all *k*-input BF classes, so it is convenient to generate EFs without any additional operations. It only depends on two steps, generating a random BF, subsequently verifying, and then accepting or regenerating. Because of EF’s trivial performance on dynamic, we omit their dynamic system analysis.

Monotonic BFs are generated with the conceptions of Boolean monotonicity. Consider two *k*-bit Boolean vectors 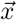 and 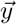, if ∀*i* ∈ {1, 2, …, *k*}, satisfies *x*_*i*_ ⩽ y_*i*_, we say 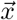 is smaller than 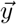, 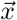 is one predecessor of 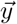, while 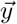 is one successor of 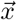. To generate a *k*-input MF, random *k* vectors are selected as predecessor seeds, followed by screening out all successors of the seeds. Subsequently, the vectors belonging to both seeds and successors are classified as set *f*^−1^(1), while their complement of entire 2^*k*^ space is set *f*^−1^(0). This approach ensures the generation of a monotonic increasing BF. Correspondingly, exchanging the categories of *f*^−1^(1) and *f*^−1^(0) returns a monotonic decreasing BF.

Post class functions stress the common component of pointed input vectors. Here we use the subset (µ = 2,{*A*^2^} ∪ {*a*^2^}) as the representative of PFs. In detail, if any two input vectors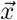, 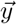 from *f*^−1^(1), ∃*i* lets *x*_*i*_ = y_*i*_ = 1, we call *f* ∈ {*A*^2^}. Inversely, {*a*^2^}.is the case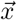, 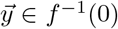, and *x*_*i*_ = y_*i*_ = 0. Especially, two vectors are *intersecting* if ∃*i* lets *x*_*i*_ = y_*i*_ = 1. To generate a *f* ∈ {*A*^2^}, we randomly select a Boolean vector 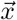 as the first element of *f*^−1^(1) (By definition,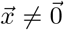. Next, select a random vector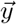, and verify whether it is intersecting with all elements of *f*^−1^(1). If it meets the criteria, add it to *f*^−1^(1); otherwise, select a new 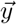. Repeat this process 2^*k*−1^ times regardless of failure or identical selections, and finally a PF∈{*A*^2^} is generated. Correspondingly, a PF∈{*a*^2^} needs above process only by replacing vectors of their bit-invert ones, and *f*^−1^(1) by *f*^−1^(0).

Threshold-based BFs have two classical types, both of which are same under σ_*i*_ ⩾ 0 but divergent in terms of σ_*i*_ < 0. One case is strictly limited to Boolean framework with binary value {0, 1}, while another exhibits a spin-like pattern (Eq.12),

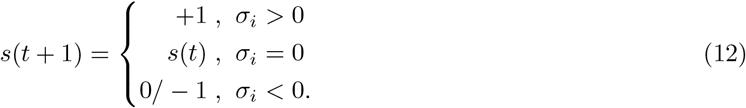

Although the spin-like mode is initially proposed for analyzing neural system [66], this setting is also utilized in modeling genetic network [31]. Here we focus solely on the Boolean mode to ensure dynamic consistency. Especially, we ignore self-loop setting in Eq. (12), namely the case σ_*i*_ = 0 returns a preassigned state, 0 or 1.

The generation can be described as follows. Firstly, randomly assign input variables belonging to positive or negative, N_(+)_ + N_(−)_ = *k*. Next, to balance positive and negative inputs, introduce another constraint that σ_*i*_ = 0 return the minority value, namely, 0 for N_(+)_ > N_(−)_, 1 for N_(+)_ < N_(−)_, and a random Boolean value for N_(+)_ = N_(−)_. Finally, fill the truth table according to Eq. (12). If limited to the order constraint, only change is fixing the value of σ_*i*_ = 0.

### Sensitivity of Boolean function and relevant energy

The definition and analysis of the sensitivity can be seen in the literature[67]. Here we directly use its formulaic expression. For a given *k*-variable SBFs, we randomly exchange any two bits of its truth table 2^*k*+1^ times as shuffled configurations.

### Hybrid Kauffman models

For simplicity, system size and in-degree of Kauffman models are fixed as 10^3^ and 4 respectively. Each parameter configuration is randomly realized 1000 times for statistics. In once implementation, a random initial state 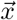 and its perturbed state 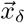 with a normal Hamming distance of 0.1, are simulated 1,000 time steps following their respective dynamic trajectories. The final distances are the averages of all realizations. The simulations and subsequent results are based on the synchronous update rule.

### Models embedded in square lattice and percolation occurrence

The method of generating a hybrid model is same as aforementioned hybrid Kauffman model. Each agent of model is positioned at the vertices of square lattice. In once simulation, we randomly embed a hybrid Kauffman model into a square lattice, and each vertex is assigned with arbitrary Boolean value. Then let systems evolve for 1000 time steps to pass transient process and to observe the stable components. A vertex that remains stationary during observing windows (500) is subsumed into stable components. It is recognized as percolation occurrence if there exists a pathway across lattices from one boundary to another.

### Biological networks and framework conversion

Biological models are collected from Cell Collective Website (Set-1, 71 models) [68] and published work (Set-2, 8 models) [32, 58, 69, 70]. We downloaded the truth tables of bio-nets of Set-1, and then analyzed their categories. For the Set-2, we use the threshold rules of each gene to establish truth tables. To identify each type of SBFs, we proposed series fast algorithms implemented by our C++ codes, which are available on GitHub. Especially, we use Z3 solver, a tool of solving the problem of Boolean satisfiability (SAT), to convert the threshold based framework to the logical one. The extended form of 𝕄 is not limit to strictly global monotonicity, namely any variables of *f* are monotonic regardless of their increasing or decreasing patterns. Extended form of 𝕋 allow weights (thresholds) to be arbitrary (nonzero) integer numbers. However, it should be noted that not all truth tables can be successfully converted to the threshold forms, namely these BFs do not belong to 𝕄_*e*_.

### Network evolution

All initial artificial networks consist of 50 nodes connected by a directed Erdős-Rényi networks. Conveniently, we treat in-/out-degrees as Poisson Distribution with λ = 5 that means connection probability is 0.1. Each node can have father or child nodes, but should not be isolated. Six operations serve as *in silico* evolution processes, namely adding a new node/edge, deleting an existing node/edge, swapping two edges, and changing the Boolean function of nodes. We assume each type share a same occurring probability due to the lack of precise existing conclusions. One operation is considered as one “generation”. It skips the “adding new nodes” operation if systems reach the maximum size set to 100. “Changing node’s Boolean function” is merely limited to replacing random BFs with CFs or TFs. Swapping edges involve exchanging two edges in the system, and its only require is to avoid duplicate connections between nodes. Simulations are terminated once systems reach predefined evolution generations, and we calculate their expected sensitivities as previously mentioned.

### Attractor domain analysis

Uncertainty arises from the use of asynchronous update rules. We assume equal priority among nodes and only one randomly selected gene can update its state in each time step. Accurately determining the periods becomes challenging when limit cycles are embedded within complex loops. Thus we categorize all complex loops into two groups according to the amount of states they contain, where threshold keeps in constant as 1,024 (= 2^10^). For one evolving-terminated network, we randomly generate a system state and let it continuously update until passing transient process (10^3^ steps). Since these networks are supposed to closing to criticality, whose lengths of transient process are acceptable in small size system. We employed a recursive method to assess the size of complex loop. Briefly, we recursively build a state transition graph of complex loops. This process is stopped until building a small state transition graph completely or state number exceeding the threshold.

For artificial evolution networks, the size of system is fixed to 50, while edges follow the Poisson distribution (λ = 5, figure 5b). 10^6^ random cases are implemented for statistics. Conversely, for real bionetworks, random states are set as min {10^7^, 2^*N*^} for testing to obtain system internal details. The actual amounts of the three types of attractors are also counted. Zero in-degree nodes are also considered as usual components though they cause systems to sink into subspace.

## Data and code availability

The source data and code for this paper are available at GitHub [71].

## Acknowledgements

This work was supported by the National Natrual Science Foundation of China (92068201, 31830060), Key R&D Program of Zhejiang (2024SSYS0031).

## Author contributions

All conceived the study. Y.Y. and Z.H. performed the theoretical analysis. Y.Y. developed and implemented the algorithms. D.P. provided biological comments and suggestions. Y.Y. wrote the draft. All revised the paper. D.P. supervised the project.

