## Supplementary Figures 1∼4 for "Canalization and competition: the cornerstone of genetic network’s dynamic stability and evolution"

### Supplementary figures for "Canalization and competition: the cornerstone of genetic network's dynamic stability and evolution"

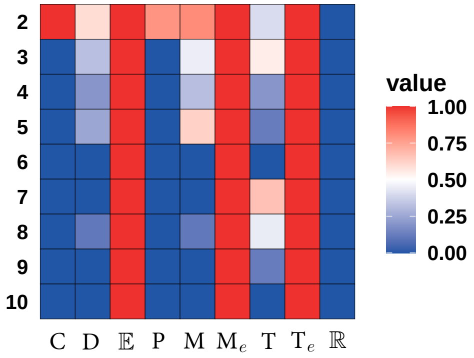


Figure S1: Abundance of SBFs in threshold based models but SBFs are converted by spin-like rules.

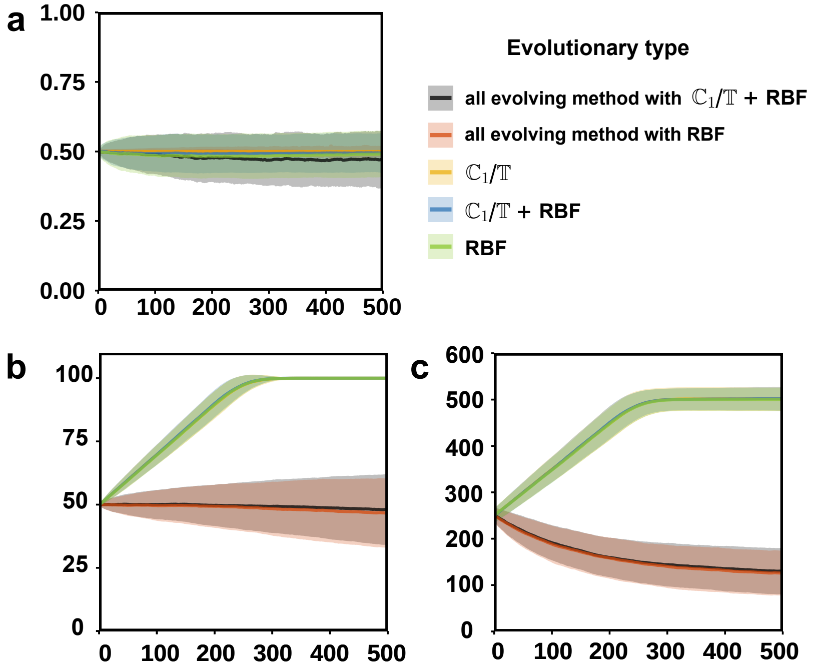


Figure S2: Bias (a), node (b), and edge (c) changing along with simulation under different evolving methods. Black and red curves are the arbitrary evolving methods as figure 3b in main text. Yellow , blue, and green curves are same as figure 3c in main text. Each curve is generated $10^3$ realizations for statistics.

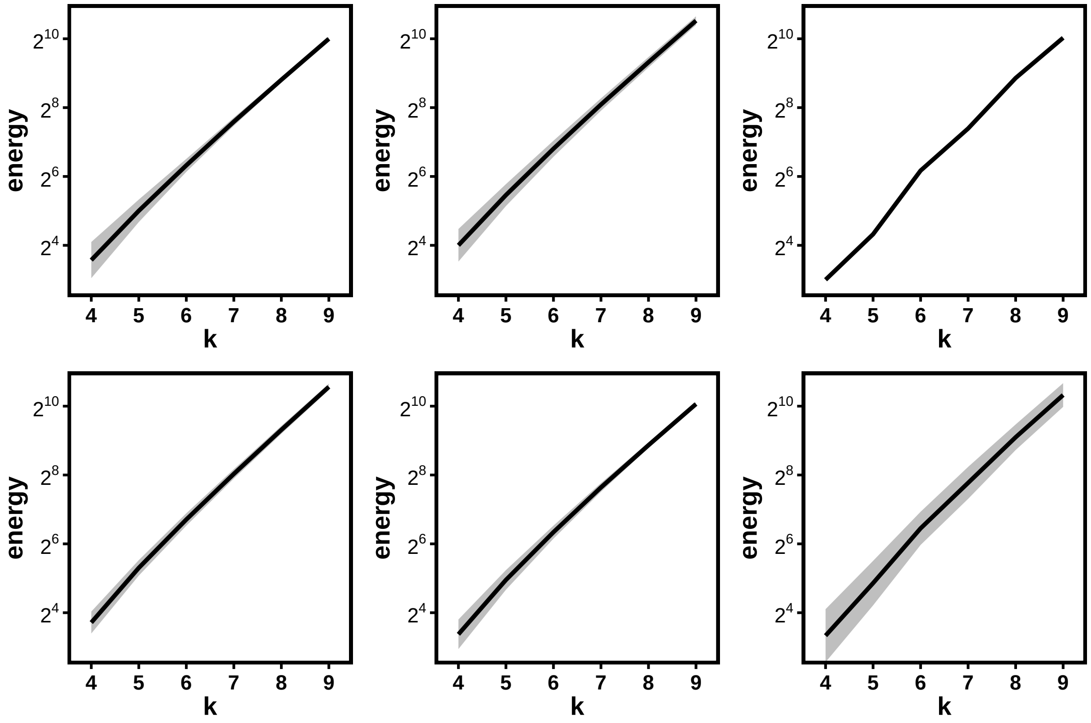


Figure S3: Pseudo energy increases exponentially as $k$ changing for detail $\mathbb{S}$ as figure 10b.

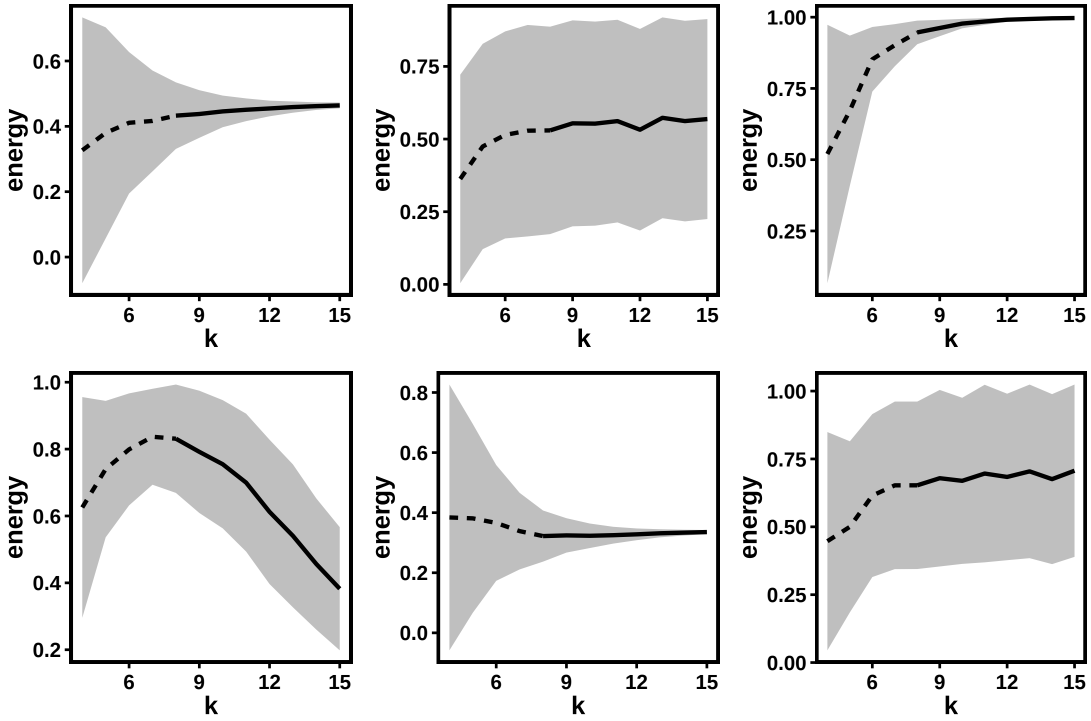


Figure S4: Relative energy of detail $\mathbb{S}$ versus $k$ as figure 10c.
